# Pannexin 1 modulates angiogenic activities of human endothelial colony-forming cells through IGF-1 mechanism and is a marker of senescence

**DOI:** 10.1101/2023.05.01.539004

**Authors:** Ting-Yi Tien, Yih-Jer Wu, Cheng-Huang Su, Chin-Ling Hsieh, Bo-Jeng Wang, Yi-Nan Lee, Yeu Su, Hung-I Yeh

**Affiliations:** Institute of Biopharmaceutical Science, National Yang Ming Chiao Tung University, Taipei, Taiwan; Department of Medical Research and; Internal Medicine, MacKay Memorial Hospital, Taipei, Taiwan; Department of Medicine, MacKay Medical College, New Taipei City, Taiwan

**Keywords:** Endothelial progenitor cells, Endothelial Colony-Forming Cells, Pannexin 1, IGF-1, FAK, calcium influx

## Abstract

**BACKGROUND:** We examined the role of pannexins in human endothelial progenitor cell (EPC) senescence.

**METHODS:** Young and replication-induced senescent endothelial colony-forming cells (ECFCs) derived from human circulating EPCs were used to examine cellular activities and senescence-associated indicators after transfection of siRNA specific to Panx1 or lentivirus-mediated Panx1 overexpression. Hindlimb ischemia mice were used as *in vivo* angiogenesis model. Protein and phospho-kinase arrays were used to determine underlying mechanisms.

**RESULTS:** Panx1 was the predominant pannexin isoform in human ECFCs and up-regulated in both replication-induced senescent ECFCs and circulating EPCs from aged mice and humans. Cellular activities of the young ECFCs were enhanced by Panx1 down-regulation, but attenuated by its up-regulation. In addition, reduction of Panx1 in the senescent ECFCs could rejuvenate cellular activities with reduced senescence-associated indicators, including senescence-associated β-galactosidase activity, p16^INK4a^, p21, acetyl-p53, and phospho-Histone H2A.X. In mouse ischemic hindlimbs injected senescent ECFCs, blood perfusion ratio, salvaged limb outcome, and capillary density were all improved by Panx1 knockdown. Insulin-like growth factor 1 (IGF-1) was significantly increased in the supernatant from senescent ECFCs after Panx1 knockdown. The enhanced activities and paracrine effects of Panx1 knockdown senescent ECFCs were completely inhibited by anti-IGF-1 antibodies. FAK, ERK and STAT3 were activated in senescent ECFCs with Panx1 knockdown, in which the intracellular calcium level was reduced, and the activation was inhibited by supplemented calcium. The increased IGF-1 in Panx1-knockdown ECFCs was abrogated respectively by inhibitors of FAK (PF562271), ERK (U0126), and STAT3 (NSC74859), and supplemented calcium.

**CONCLUSIONS:** Panx1 expression is up-regulated in human ECFCs/EPCs with replication-induced senescence and during aging. Angiogenic potential of senescent ECFCs is improved by Panx1 reduction through increased IGF-1 production via activation of FAK-ERK axis following calcium influx reduction. Our findings provide new strategies to evaluate EPC activities and rejuvenate senescent EPCs for therapeutic angiogenesis.

## Introduction

Endothelial progenitor cells (EPCs) are capable of re-endothelialization in injured vessels and neovascularization in ischemic tissues.^1^ Chemoattractants such as stromal cell-derived factor 1 (SDF-1) or stem cell factor (SCF) released from injured endothelial tissue play an important role in stimulating differentiation and migration of EPCs. The number and function of EPCs declined during the process of senescence induced by excess oxidative stress and inflammation, or decreased pro-angiogenic factors, e.g., insulin-like growth factor 1 (IGF-1), which reduces senescence through FAK-ERK axis and enhanced eNOS and VEGF production in EPCs.^2, 3^ Senescence of EPCs may lead to endothelial dysfunction followed by cardiovascular diseases.^4^ Autologous transplantation of EPCs have been applied for clinical trials of ischemic disease, such as acute myocardial infarction and peripheral arterial disease, and showed improved outcomes.^5, 6^ One of the challenges in application of EPCs for therapy is that patients with cardiovascular diseases are commonly old, and EPCs from these patients usually present impaired cellular activities due to senescence, leading to poor therapeutic outcomes.^7^ In addition, for therapeutic angiogenesis, an adequate number of injected cells is necessary to achieve the effect.

Endothelial colony-forming cells (ECFCs), one population of EPCs possessing proliferation activities and endothelial angiogenic potential, could be isolated form peripheral blood and cultured to several passages, and were considered as real EPCs. ECFCs have been reported to apply to tissue engineering and clinical trials of ischemic disease.^8^ However, along the passages of ECFCs, replication-induced senescence attenuates the angiogenic activities. These difficulties highlight an unmet need for a new marker to reflect activities and senescence of EPCs/ECFCs and to develop strategies to improve therapeutic potential of senescent EPCs/ECFCs.

Pannexins are mammalian transmembrane proteins possessing sequence homology to innexins and topologic homology to connexins, gap junction proteins in invertebrate and vertebrate, respectively. Unlike innexins and connexins, pannexin molecules could be glycosylated at extracellular loops.^9^ There are three pannexin isoforms (Panx1, Panx2, and Panx3) that form homomeric conductance channels in cell membrane, named pannexons, which allow the passage of molecules smaller than 1 kDa, such as ATP and Ca^2+^, through cell membrane. Recent evidences from cryogenic electron microscopy showed that each pannexon channel can be constituted by different number of pannexin isoforms.^10, 11^ Among the three pannexin isoforms, Panx1 is widely distributed in many cell types and tissues, including vascular endothelium, while Panx2 and Panx3 are restrictively expressed.^12^ Panx1 channels are involved in inflammasome formation and apoptopsis.^13, 14^ In cardiovascular system, the studies on Panx1 systemic and renin expressing cell knockout mice showed that Panx1 maintains electrophysiological function and blood pressure.^15, 16^ Other studies in endothelial cells showed that TNF-α may stimulate Panx1 production through activation of NFκB pathway and trigger ATP release from pannexons through activation of Src family kinase (SFK), leading to neutrophil adhesion onto endothelium.^17, 18^ In keratinocytes, reactive oxygen species (ROS) was also reported to induce opening of pannexin channels.^19^ These findings indicated that the expression and function of Panx1 could be influenced by inflammatory cytokines and oxidative stress.

Our previous works on the role of connexin43 (Cx43), the predominant connexin isoform in ECFCs, showed that reduction of Cx43 impaired their cellular activities and therapeutic angiogenic potential of smooth muscle progenitor cells through FAK activation, indicating maintenance of progenitor cell function requires Cx43.^20, 21^ However, the roles of pannexins in EPCs/ECFCs remained unclear. We hypothesized that Panx1 expression in EPCs can be influenced by inflammatory cytokines and oxidative stress elevated in senescent process of subjects, and the change may be related to cellular activities, including senescence, of EPCs/ECFCs.^22^

In this study, we used ECFCs for *in vitro* experiments and established a senescent model. To dissect the role of Panx1 on angiogenic potential of human ECFCs, cellular activities of young ECFCs after perturbation of Panx1 using short interfering RNAs (siRNAs) and lentivirus–mediated Panx1 overexpression were analyzed. To clarify whether Panx1 is involved in modulating senescence process of human ECFCs, we assessed cellular activities and senescent-associated indicators of the senescent ECFCs after Panx1 reduction *in vitro* and in mouse hindlimb ischemia model. In addition, the underlying mechanisms were also explored.

## Methods

### Data Availability

The supplemental data and study materials related to this study are available to other researchers on reasonable request. Methods are expanded in the <u>Supplemental Material</u>.

## Results

### Characterization and establishment of replication-induced senescence modle in human ECFCs

CD34^+^ cells were isolated from peripheral blood mononuclear cells (PBMNCs) of young (20 to 30 years old) healthy donors using magnetic beads conjugated with anti-CD34 antibodies and then cultured in MV2 medium. After 14 days, cobble-stone like colonies, also called ECFCs, appeared and were passaged for subsequent experiments (Figure S1A).^16^ Eight ECFCs strains form these colonies growing to passage 16 or more were used in the *in vitro* experiments.^21^ To obtain enough cell number for experiment, ECFCs were characterized at passage 4 by flow cytometry. CD45 and CD14 (leukocyte markers) were not expressed on these cells, but CD34, KDR, AC133, and CD31 (considered as ECFC markers) could be found and the expression profiles resembled human microvascular endothelial cells (HMVECs) (Figure S1B and S1C). Surface marker expression in the 8 ECFC strains applied in this study is listed in supplemented table (Table S1). In order to evaluate the distribution of pannexin isoforms in the ECFCs, real-time PCR was conducted and the results showed that the predominant pannexin isoform of the cells was Panx1, followed by Panx2 (about 1/100 of Panx1), while Panx3 was virtually absent (Figure S2A).

To compare young and senescent ECFCs in Panx1 expression (Figure 1), senescence was induced by replication. We defined passages 6 to 8 of ECFCs as young group and passages 14 to 16 as senescent group and found that doubling time of senescent ECFCs was more than 3 folds longer than that of young ECFCs (senescent vs. young/ 106.5±19.1 vs. 28.3±5.2 hours)^23^. In addition, telomere length (telomere repeat copy number (T)/ single-copy gene 36B4 copy number (S) ratio) of senescent ECFCs was about 40% shorter, compared to young ECFCs, similar to previous reports (Figure S2B).^23, 24^ In parallel, compared to young ECFCs, the expression of senescence-associated indicators in senescent ECFCs, including p16^INK4a^, p21, acetyl-p53, and senescence-associated beta-galactosidase activity (SA-β-gal) were also enhanced (Figure S2C and S2D). In subsequent experiments, phospho-Histone H2A.X (phosphor-H2A.X, a DNA damage marker), manganese superoxide dismutase (MnSOD), and IL-6, were also enhanced in senescent ECFCs, while SirT1 and Lamin B1 were decreased but glutathione peroxidase (GPx) remained steady. Cellular activities of proliferation, tube formation, and migration of the ECFCs at passages 14 to16 were about half to one fifth less, compared to the cells at passages 6 to 8 (Figures 2 and 3).

**Figure 1.**
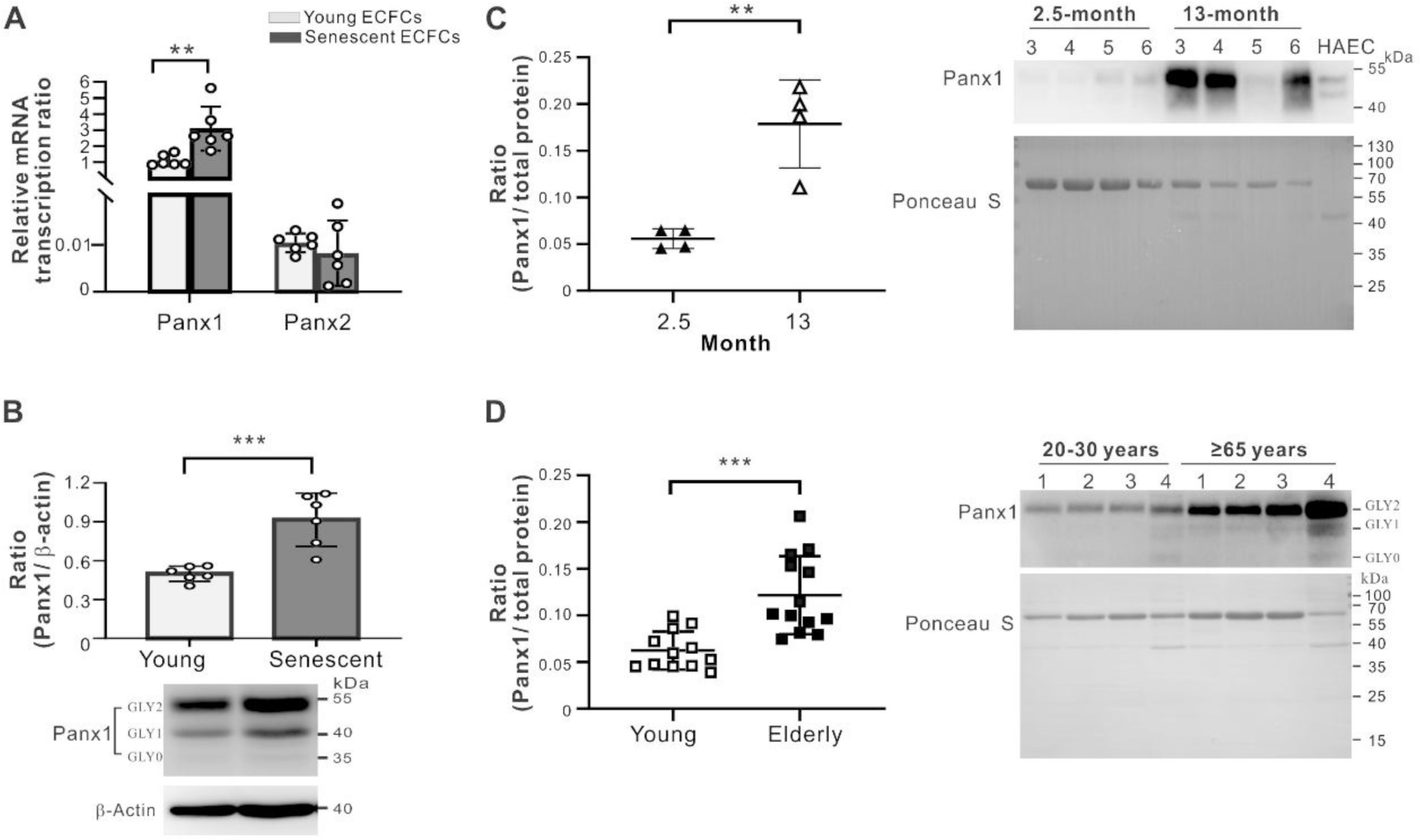
Enhanced Panx1 expression in senescent ECFCs and circulating EPCs obtained from aged mice and humans. **A and B,** Expression of pannexin isoforms was analyzed in young (passages 6 to 8) and senescent (passages 14 to 16) ECFCs. Real-time PCR showed that transcript expression of Panx1 in senescent ECFCs was twice more than that in young ECFCs, while Panx2 expression remained steady (**A**). Western blotting confirmed increased Panx1 protein expression in senescent ECFCs (**B).** N=6, individual donors. **C,** Comparison of Panx1 expression in circulating EPCs (Scal-1^+^) between 2.5 and 13-month old mice (n=4 in each group) by western blotting**. D,** Comparison of Panx1 expression in circulating EPCs (CD34^+^) between young (20 to 30 years old, n=12) and elderly (**≥** 65 years, n=13) donors by western blotting. Note that protein expression levels of Panx1 in circulating EPCs are higher in aged mice and people, compared to the young corresponding groups. Data are mean ± SD. Statistical significance of all results was assessed by unpaired t-test. *, P<0.05. **, P<0.01. ***, P<0.001.

**Figure 2.**
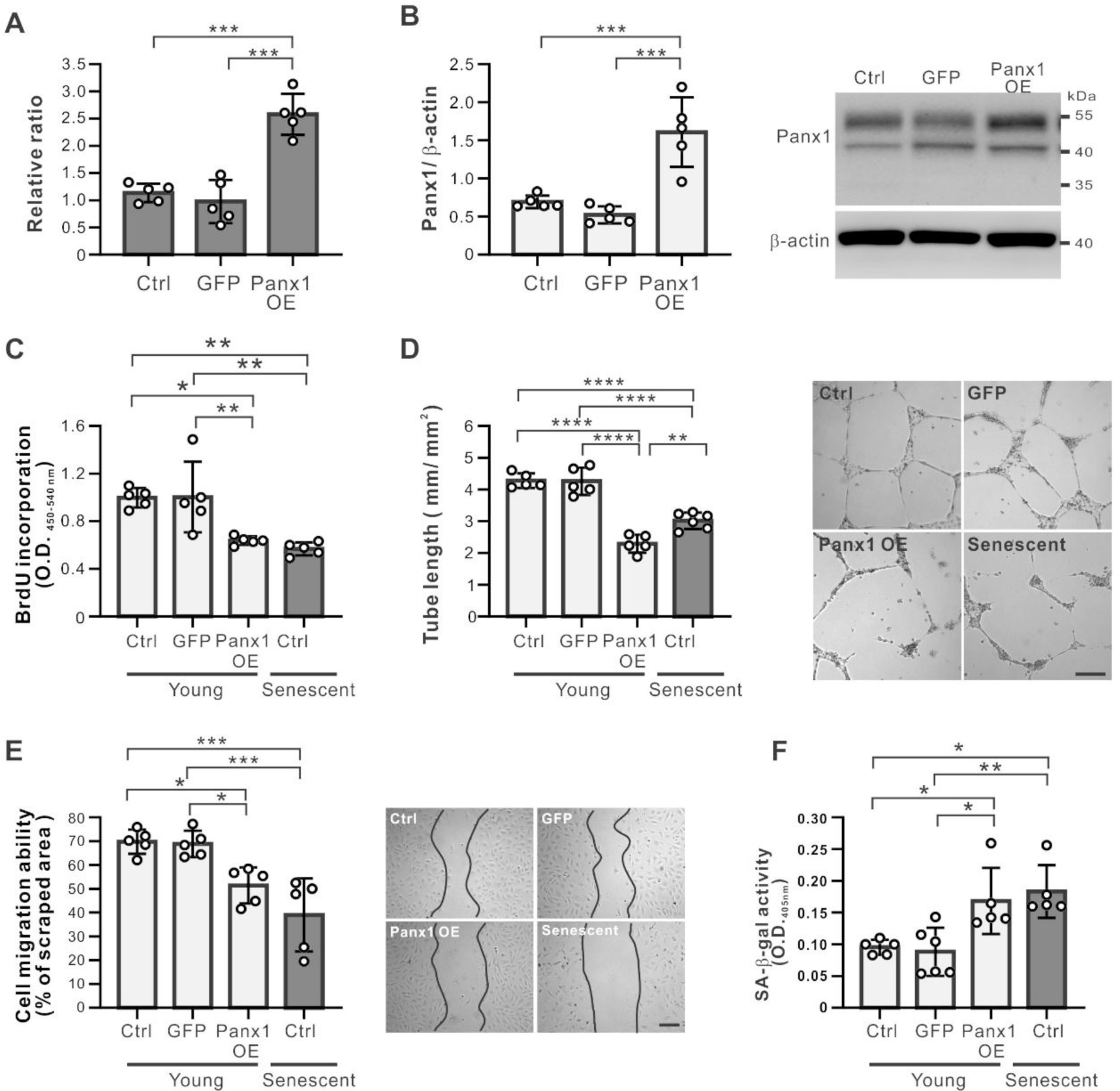
Cellular activities of young ECFCs are impaired by Panx1 overexpression. **A and B,** 2 to 3-fold overexpression of Panx1 in young ECFCs was achieved post infection with lentivirus carrying Panx1 cDNA fragments, as detected by real-time PCR (**A**) and western blotting (**B)**. **C-E,** Young ECFCs with overexpressed Panx1 had attenuated activities of proliferation (**C**), tube formation (**D**), and migration (**E)**. Bars, 200 μm in **D** and **E**. **F**, Measurement of SA-β-gal activity (optical density measurement, 405 nm) in young ECFCs showed that the activity was enhanced by Panx1 overexpression, nearly to that in senescent ECFCs. N=5 to 6 ECFC strains from individual donors in each assay. Data are mean ± SD. Statistical analysis was performed by one-way ANOVA followed by the Tukey’s test. *, P<0.05. **, P<0.01. ***, P<0.001, ****, P<0.0001.

**Figure 3.**
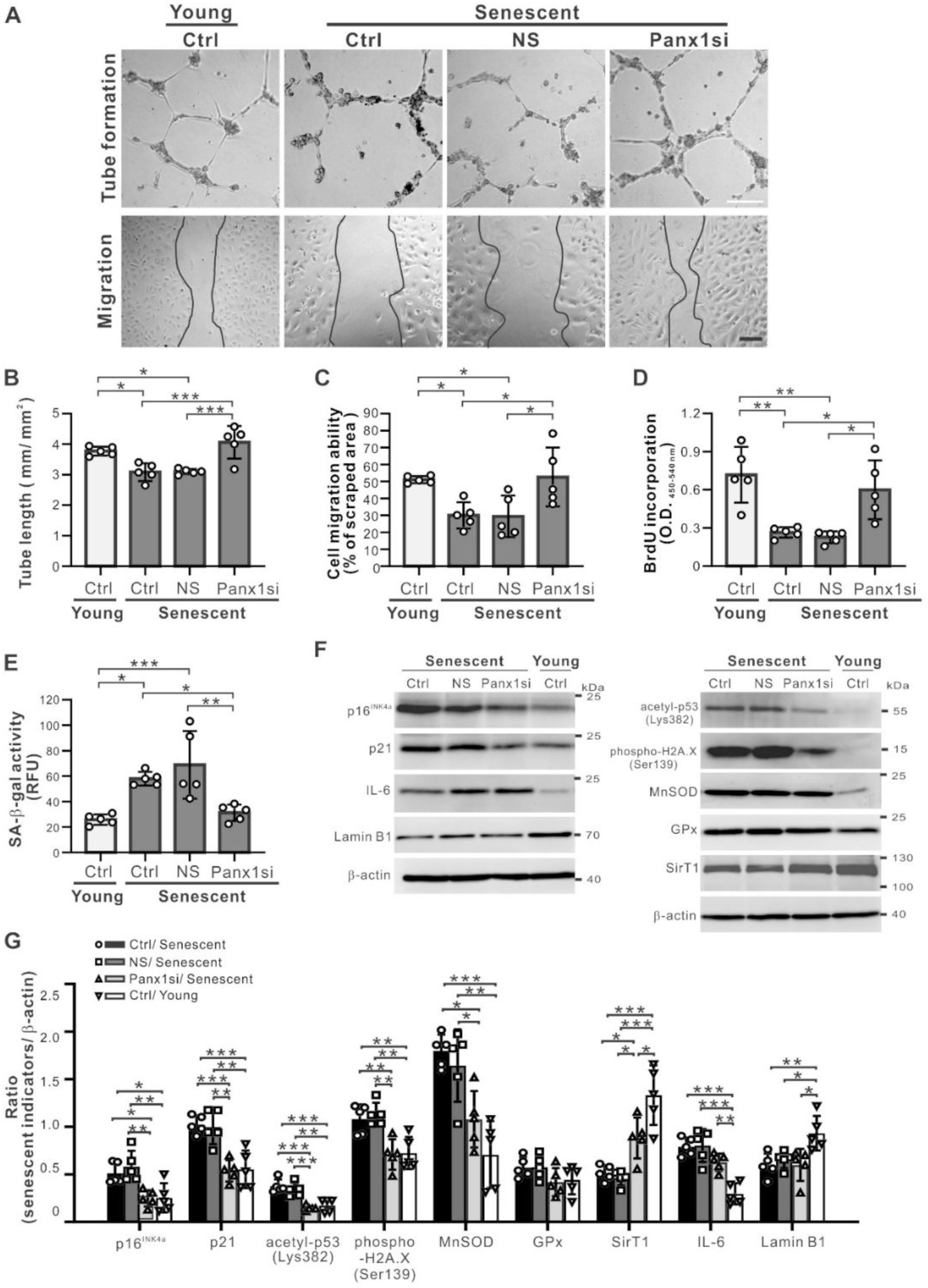
Panx1 knockdown enhances cellular activities and reduces the expression of senescence-associated markers in senescent ECFCs. **A-D,** Evaluation of cellular activities of senescent ECFCs showed that tube formation (**A, upper panel, and B**), migration (**A, lower panel, and C**), and proliferation (**D**) were enhanced to near young ECFCs levels at 24 hours post Panx1 siRNA transfection. Bars: 200 μm in **A**. **E,** Measurement of senescence-associated β-galactosidase (SA-β-gal) activity in Panx1 knockdown senescent ECFCs (fluorescence measurement, 490/510 nm) showed that the activity in senescent ECFCs was reduced to near young ECFC level. **F and G,** Western blotting images (**F**) and density statistics (**G**) show that, in senescent ECFCs after Panx1 knockdown, p16^ink4a^, p21, acetyl-p53, phospho-H2A.X and MnSOD were reduced and Sirt1 was increased, while GPx, IL6 and lamin B1 remain steady. N=5 ECFC strains from individual donors in each assay. Data are mean ± SD. Statistical analysis was performed by one-way ANOVA followed by the Tukey’s test. *, P<0.05. **, P<0.01. ***, P<0.001.

### Enhanced Pannexin 1 expression in senescnet ECFCs and circulating EPCs of aged mice and humans

The expression of pannexin isoforms was compared between young and senescent ECFCs. The transcript and protein expression levels of Panx1 were more than 90% higher in the senescent ECFCs, while the transcripts of Panx2 remained stationary, compared to the young cells (Figure 1A and 1B), indicating that expression of Panx1 was sensitive to the senescence status of ECFCs. Immunoconfocal images also confirmed the enhanced expression of Panx1 in cell membrane and cytoplasm of senescent ECFCs (Figure S3A). In order to verify the enhanced Panx1 expression associated with senescence *in vitro* also existed *in vivo*, mouse circulating EPCs (Scal-1^+^ PBMNCs) were isolated from young (2.5-month) and old (13-month) animals. Western blotting showed that Panx1 expression level in circulating EPCs from old mice was nearly 3-fold higher, compared to young mice (Figure 1C). Moreover, human circulating EPCs (CD34^+^ PBMNCs) were isolated from 12 young (20 to 30 years, 5 males) and 13 elderly (≥65 years, 6 males) donors. Consistent with the mouse findings, Panx1 expression in circulating EPCs from elderly donors was enhanced about 2-fold than that from young donors (Figure 1D and Figure S4B). Biochemical examination of the donors revealed that fasting plasma glucose (FPG), triglycerides (TG), glutamate pyruvate transaminase (GPT), glutamic-oxaloacetic transaminase (GOT), and blood urea nitrogen (BUN) were higher in the elderly donors (Table S2). These results indicated that Panx1 expression was up-regulated when the ECECs/EPCs become senescent both *in vitro* and in *vivo*, suggesting that Panx1 expression may affect cell function during the senescence process.

### Pannexin 1 expression level determines cellular activities of ECFCs

To clarify the role of Panx1 in biological function of ECFCs, Panx1 expression was respectively up-regulated and down-regulated in young ECFCs using lentiviruses carrying Panx1 cDNA fragments and two different siRNA (Panx1si-A and Panx1si-B) specific to Panx1 and followed by cellular activity evaluation. In young ECFCs treated with the lentiviruses carrying Panx1 cDNA fragments, the expression of Panx1 mRNA and protein were increased about 2.6-fold, similar to the level of senescent ECFCs (Figure 2A and 2B). Immunoconfocal images confirmed the enhancement in both cell membrane and cytoplasm (Figure S3A). To evaluate the behaviors of Panx1 up-regulated ECFCs, BrdU incorporation, tube formation, and wound healing assay were conducted. The results showed that proliferation, tube formation and migration of young ECFCs could be decreased by Panx1 up-regulation (Figure 2C to 2E). In addition, SA-β-gal activity in young ECFCs could also be increased by Panx-1 up-regulation (Figure S3B) nearly to the level of senescent ECFCs (Figure 2F). These results indicated that increased Panx1 expression impaired cellular activities of young ECFCs.

In Panx1 siRNA transfected young ECFCs, the mRNA expression could be reduced about 85 to 95% by each of the two siRNA and the reduction of Panx1 did not cause compensatory up-regulation of Panx2 (Figure S5A). Western blotting showed that Panx1 expression of ECFCs was also reduced about 80% by each of the two siRNAs and immunoconfocal images confirmed the reduction in both cell membrane and cytoplasm (Figure S5B and S5C). As expectedly, proliferation, tube formation and migration of young ECFCs could be enhanced by Panx1 reduction (Figure S5D to S5F).

Collectively, these findings revealed that Panx1 not only comprised pannexon channels but the expression was also closely related to cellular activities of ECFCs. Since Panx1 expression was enhanced in senescent ECFCs and down-regulation of Panx1 increased cellular activities of young ECFCs, these findings suggested that reduction of Panx1 in senescent ECFCs may rejuvenate the cellular function. Due to the effects of Panx1si-A and Panx1si-B were compatible, Panx1si-A was used in subsequent experiments focusing on senescent ECFCs.

### Reduction of Panx1 improves angiogenic potential of senescent ECFCs *in vitro and in vivo*

To examine the hypothesis that Panx1 reduction could rejuvenate angiogenic potential of senescent ECFCs, tube formation (Figure 3A, upper panel and 3B), migration (Figure 3A, lower panel and 3C), and proliferation assay (Figure 3D) were conducted in senescent ECFCs after transfection of Panx1 siRNA-A. The results showed that the three activities of Panx1 knockdown senescent ECFCs could be augmented near to or even over the levels of young ECFCs. In order to understand if Panx1 was involved in senescence of ECFCs, SA-β-gal activity and senescent associated indicators were detected in Panx1-down-regulated senescent ECFCs. SA-β-gal activity assay and immunoconfocal images showed that the enhancement of SA-β-gal activity, and phospho-Histone H2A.X in senescent ECFCs could be reduced by Panx1 reduction (Figure 3E and Figure S6). Moreover, the enhancement of p16^INK4a^, p21, acetyl-p53, phospho-Histone H2A.X, and MnSOD in senescent ECFCs was reduced and the decrease of SirT1 was increased, but GPx, IL-6, and Lamin B1 were little affected after Panx1 reduction (Figure 3F and 3G). These findings indicated that Panx1 reduction not only rejuvenate cellular activity but also reversed senescent status of ECFCs.

Endothelial nitric oxide synthase (eNOS), an important factor modulating ECFC function, was also examined. After Panx1 knockdown, the decrease of phosphorylated eNOS (Ser1177) and total eNOS in senescent ECFCs were insignificantly increased and remained significantly lower than that of young ECFCs (Figure S7A to S7C). In parallel, compared to virus untreated (control) group or GFP bearing-lentivirus transfected group, the expression of total eNOS was decreased in Panx1 overexpressed young EPCs, but the ratio of phosphorylated eNOS was steady (Figure S7D to S7F). These results indicated that ECFC dysfunction induced by Panx1 overexpression involved decrease of total eNOS, but cellular activities of senescent ECFCs enhanced by Panx1 reduction was through factors other than eNOS.

To further clarify whether therapeutic angiogenesis using senescent ECFCs could be improved by Panx1 reduction, mouse hindlimb ischemia model was applied and senescent ECFCs after non-sense or Panx1 siRNA transfection were injected into the ischemic hindlimbs. Afterwards, blood perfusion ratio (blood perfusion of ischemic hindlimb divided by normal hindlimb) was recorded using Laser Doppler Imager from day 0 to day 21 and analyzed. Amputation, toe necrosis, and salvage outcome of the hindlimbs were also evaluated at the end of experiment. The results (Figure 4A and 4B) showed that blood perfusion ratios in mice injected with PBS, non-transfected (control group) and non-sense (NS) siRNA transfected senescent ECFCs were lower than that of young ECFC treatment group. However, the ratio in mice treated with Panx1 knockdown ECFCs was enhanced, nearly to the level of young ECFCs from day 14 to day 21 post injection (Figure 4B and 4C). In addition, regarding appearance of the ligated hindlimbs at the end of experiment, Panx1 knockdown senescent ECFCs injection group was free from amputation and toe necrosis, one or both of which existed in the other groups (Figure 4D). Furthermore, ischemic hindlimb tissue stained with bs-1 lectin and anti-laminin antibodies showed that the capillary density in Panx1 siRNA transfected senescent ECFC group was comparable to young ECFC group and higher than the other groups (Figure 4E and 4F). These findings indicated that Panx1 reduction may recover angiogenic potential of ECFCs impaired by senescence.

**Figure 4.**
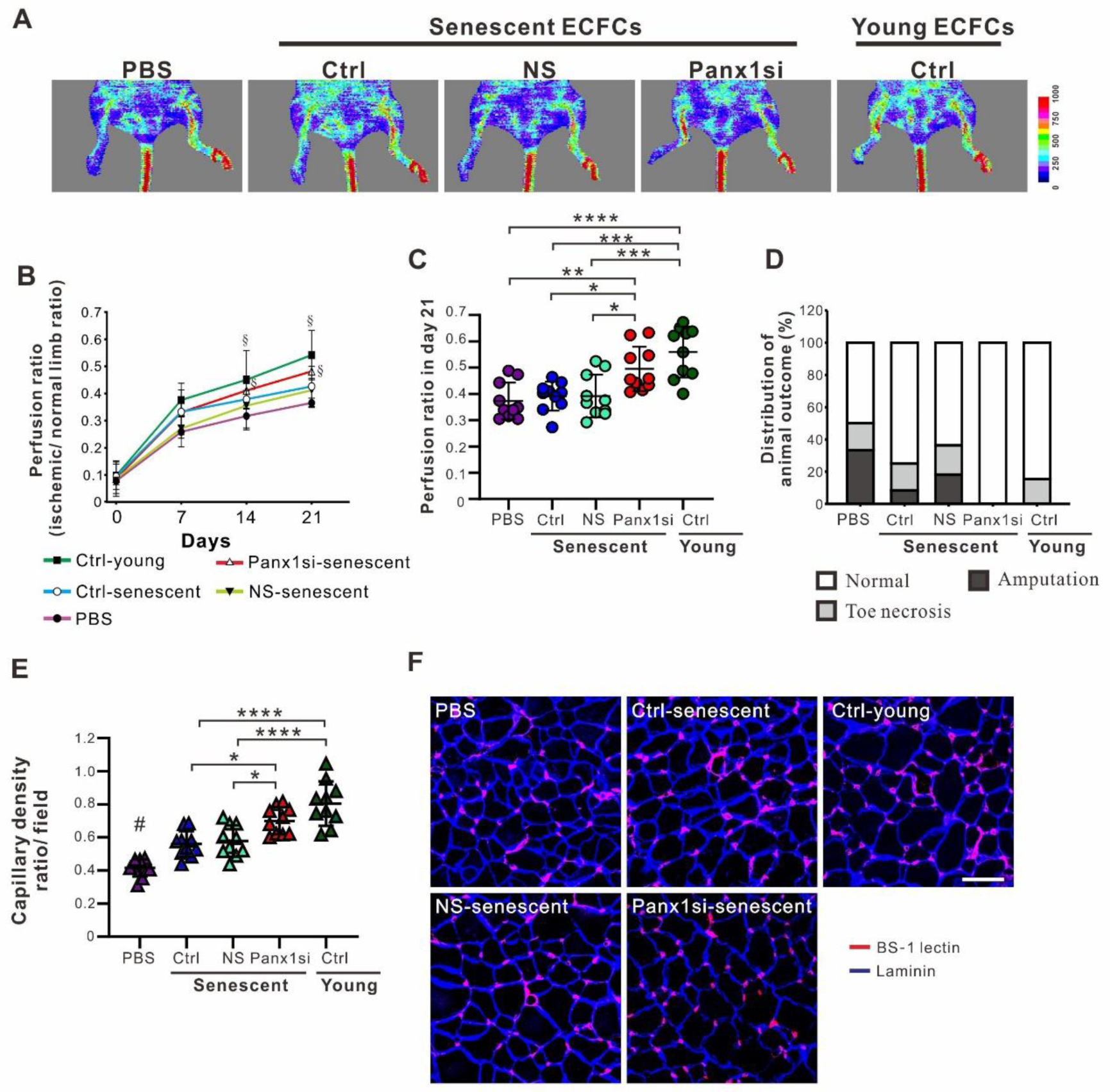
Therapeutic angiogenesis of senescent ECFCs in mouse ischemic hindlimb is improved by Panx1 reduction. **A-C,** Blood perfusion ratio in ischemic hindlimbs of mice was recorded by laser Doppler perfusion imaging from day 0 to day 21 after injection of senescent ECFCs with or without transfection of Panx1siRNA. Representative images (day 21, **A**) and statistic graph (**B**) show that the perfusion ratios of groups having ligated hindlimbs injected with young ECFCs or Panx1siRNA transfected senescent ECFCs are higher, compared to nonsense siRNA (NS) transfected senescent ECFC group, senescent ECFCs without transfection (Ctrl) group, and PBS group. The perfusion ratios between young ECFC group and Panx1 siRNA transfected senescent ECFC group are indiscriminate at day 14 and day 21 (**B and C**). **D**, Analysis of limb outcome showed that normal limb rate of Panx1 knockdown senescent ECFC group is higher, compared to PBS, Ctrl or NS senescent ECFC group, while similar to the rate of young ECFC group. **E and F**, Evaluation of capillary density by the number of capillaries (bs-1 lectin, red) divided by the number of muscle bundles (laminin signal, blue) in calf of the mice showed results consistent with the data of **C**. N=10 for each group. Bar, 50 μm. Data are mean ± SD. Statistical analysis was performed by one-way ANOVA followed by the Tukey’s test. *, P<0.05. **, P<0.01. ***, P<0.001. ****, P<0.0001. §, P<0.05, compared to PBS, Ctrl-senescent, and NS-senescent group. #, P<0.05, compared to each of the other groups.

Collectively, these consistent findings revealed that Panx1 reduction rejuvenated senescent ECFCs *in vitro* and lead to improved therapeutic angiogenesis *in vivo*.

### Improved angiogenic potential of senescent ECFCs by Panx1 knockdown is mainly attributed to insulin-like growth factor 1

Secretion of growth factors and cytokines of ECFCs has been shown to contribute to their angiogenic potential.^20, 21^ To clarify if autocrine and/or paracrine effects are involved in improved angiogenic potential of ECFCs after Panx1 knockdown, supernatants between those from non-sense transfected ECFCs and those from Panx1 siRNA transfected ECFCs were compared using protein array assay (Figure 5A). The results showed that insulin-like growth factor 1 (IGF-1) was higher and IL-1α and CCL5 were lower in amount in Panx1 knockdown senescent ECFCs versus nonsense siRNA transfected ECFCs (Figure S8A). The finding of IGF-1 was further confirmed by ELISA (Figure 5B), while the changes of IL-1α and CCL5 were inconspicuous (Figure S8B and S8C). In addition, IGF-1 in supernatants of young ECFCs was also higher in amount, compared to senescent ECFCs (Figure 5C). These results suggested that IGF-1 production was reduced during senescence process and Panx1 reduction promoted IGF-1 production in senescent ECFCs.

**Figure 5.**
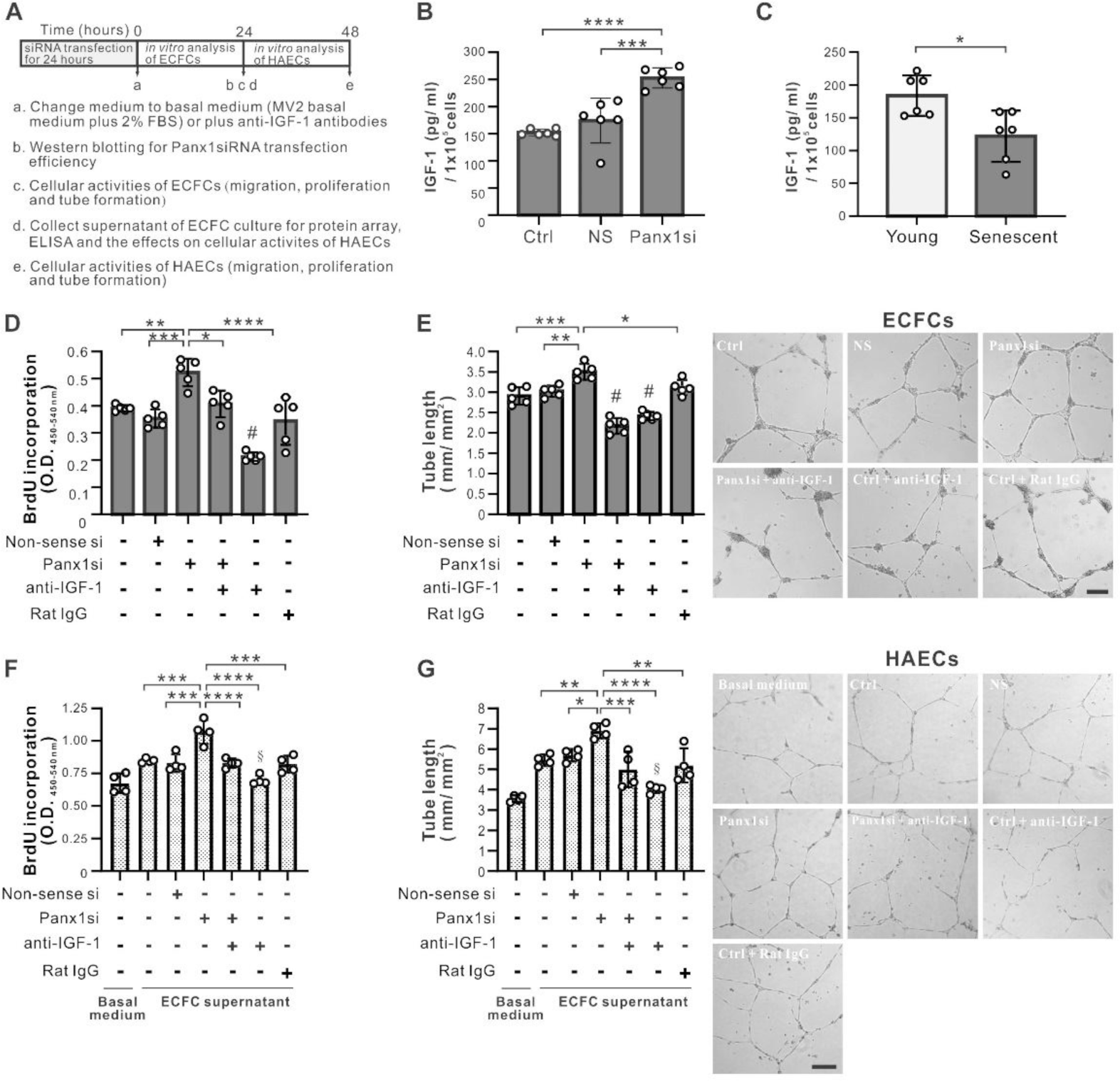
Angiogenic potential and autocrine/paracrine effects of senescent ECFCs are improved by Panx1 reduction through IGF-1. **A,** Schematic figure of experiment design and time courses. **B,** ELISA showed enhanced IGF-1 production from senescent ECFCs (passages 13 to 14) post Panx1 down-regulation (n=6, individual donors). **C,** ELISA showed lower IGF-1 production from senescent ECFCs, compared to young ECFCs (n=6, individual donors). **D and E,** BrdU assay (**D**) and matrix-gel culture (**E**) showed proliferation and tube formation were enhanced in Panx1 siRNA-transfected senescent ECFCs and the enhancement was attenuated by addition of anti-IGF-1 antibodies. N=5, individual donors. #, P<0.05, compared to each of the other bars without # above. **F and G,** BrdU assay (**F**) and matrix-gel culture (**G**) showed proliferation and tube formation of HAECs were enhanced post treatment with supernatants from Panx1 siRNA-transfected ECFCs and the enhancement was attenuated by IGF-1 neutralization. N=4, individual donors. §, P<0.05, compared to each of the other groups, except basal medium group. Bars in E and G, 200 μm. Data are mean ± SD. Statistical analysis was performed by one-way ANOVA followed by the Tukey’s test. *, P<0.05. **, P<0.01. ***, P<0.001. ****, P<0.0001.

To verify that IGF-1 was involved in the change of cellular activities of ECFCs in response to Panx1 reduction, neutralizing antibodies against IGF-1 were added to Panx-1 siRNA transfected senescent ECFC culture. Such a treatment with anti-IGF-1 antibodies abolished the improved activities of proliferation, tube formation, and migration of senescent ECFCs seen after Panx1 reduction (Figure 5D and 5E, Figure S9A), indicating that IGF-1 played a key role to enhance cellular activities of senescent ECFCs with Panx1 knockdown.

Besides, to evaluate the contribution of IGF-1 on the autocrine/paracrine effects of Panx1 knockdown senescent ECFCs, cellular activities of human aortic endothelial cells (HAECs) were analyzed after addition of supernatants from senescent ECFCs, nonsense siRNA transfected senescent ECFCs, and Panx1 siRNA transfected senescent ECFCs, with and without anti-IGF-1 antibodies. The results showed that activities of proliferation, tube formation, and migration of HAECs were improved by supernatants from senescent ECFCs and the improvement was enhanced by Panx1 reduction in senescent ECFCs, which was attenuated by addition of anti-IGF-1 antibodies (Figure 5F and 5G, Figure S9B). These results indicated that better paracrine effects of senescent ECFCs on cellular activities of HAECs by Panx1 knockdown of ECFCs was mainly through IGF-1.

### Increased IGF-1 production from ECFCs by Panx1 reduction is through activation of FAK-ERK axis

To clarify the mechanisms underlying increased IGF-1 production from Panx1 knockdown ECFCs, activation of signal transduction pathway was screened using phospho-kinase array. The results showed that phosphorylation of ERK (T202/Y204) and STAT3 (S727) was enhanced in Panx1 knockdown ECFCs (Figure S10). Previous studies reported that IGF-1 production was related to activation of ERK or STAT3, and STAT3 could be phosphorylated by ERK at the site of serine 727.^25–27^ In addition, production of cytokines and growth factors mediated via FAK-ERK axis and, in parallel, activation of FAK by channel protein reduction, such as connexin43 or P2X7 receptor, have also been reported.^21, 28, 29^ Based on these evidences, we suspected that increased IGF-1 production by Panx1 knockdown may be through activation of FAK-ERK-STAT3 axis. To verify the hypothesis, phosphorylation of FAK, ERK, and STAT3 was detected in senescent ECFCs at 30 minutes post 24-hour non-sense or Panx1 siRNA transfection and the results showed that the phosphorylation of FAK, ERK, and STAT3 were enhanced in ECFCs with Panx1 knockdown and the enhancement was attenuated by addition of FAK inhibitor (PF562271), indicating that activation of ERK and STAT3 came from activated FAK (Figure 6B to 6E). In addition, IGF-1 was measured in supernatants from Panx1-down-regulated ECFCs treated with or without ERK inhibitor (U0126), STAT3 inhibitor (NSC74859), or FAK inhibitor (PF562271) and the results showed that each of the three inhibitors could abolish IGF-1 production increased by Panx1 knockdown (Figure 6F). These findings indicated that IGF-1 production enhanced by Panx1 knockdown in ECFCs was mainly through FAK-ERK-STAT3 axis.

**Figure 6.**
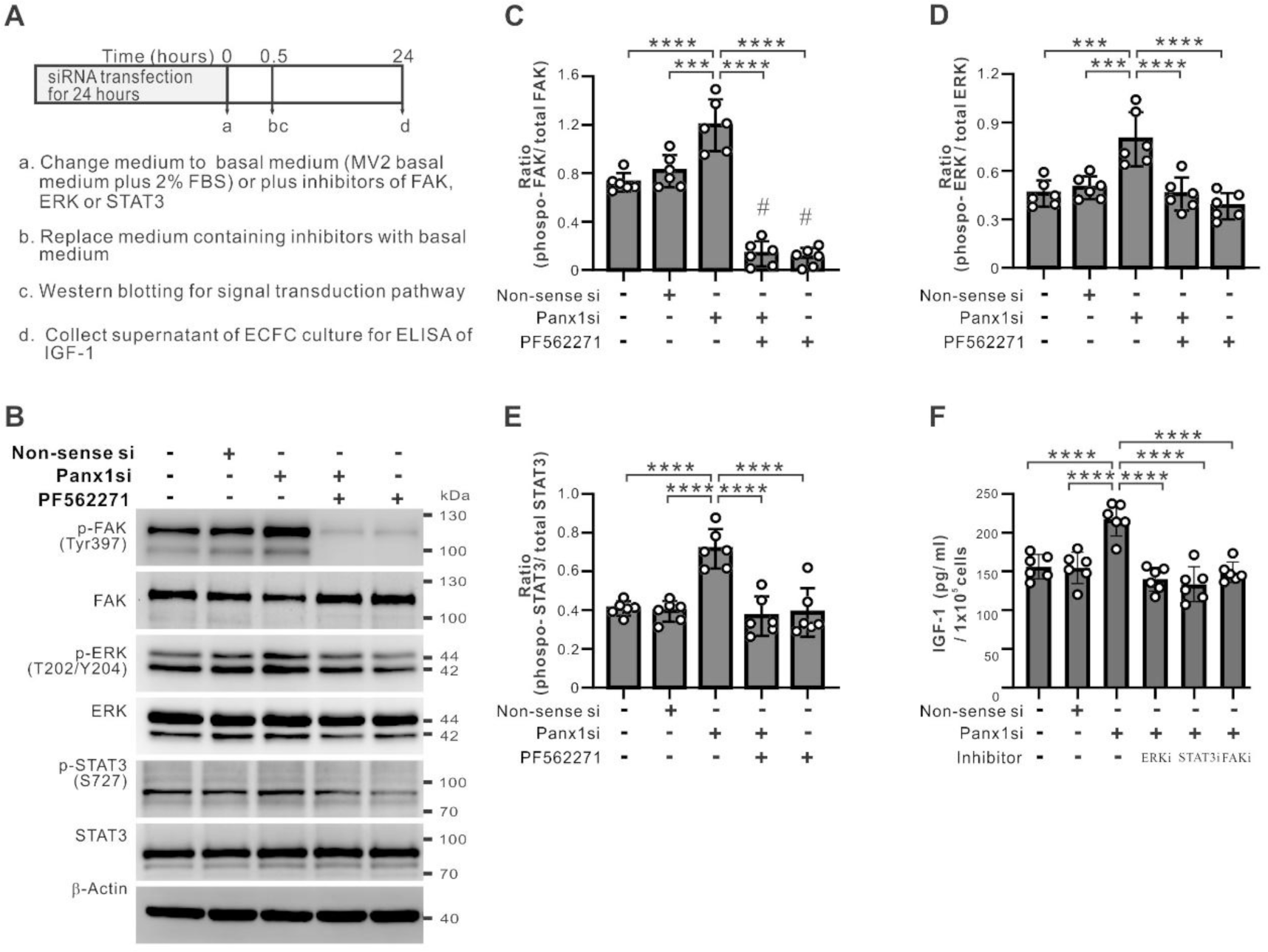
Activation of FAK-ERK axis in senescent ECFCs with Panx1 knockdown enhances IGF-1 production. **A,** Schematic figure of experiment design and time courses of inhibitor treatment. **B-E,** Western blotting (**B**) and density statistics showed that the phosphorylation of FAK (**C**), ERK (**D**) and STAT3 (**E**) in ECFCs was enhanced by Panx1 knockdown at 30 minutes post siRNA transfection and the enhancement was inhibited by addition of FAK inhibitor (PF562271, 10 μM). **F**, ELISA showed enhanced IGF-1 production from senescent ECFCs after Panx1 siRNA transfection was attenuated by addition of each of ERK inhibitor (U0126, 10 μM), STAT3 inhibitor (NSC74859, 10 μM), and FAK inhibitor (PF562271, 10 μM). N=6 ECFC strains from individual donors. Data are mean ± SD. Statistical analysis was performed by one-way ANOVA followed by the Tukey’s test. ***, P<0.001. ****, P<0.0001. #, P<0.05, compared to each of the other bars without # above.

### FAK activation in Panx1-down-regulated ECFCs resulted from decreased calcium influx

As previous studies of connexin43 and P2X7 receptor and in the present study of Panx1, we found FAK activation was involved in reduction of all 3 channels. Common small molecules pass through the three channels were compared and involvement of calcium influx was suspected.^18, 30, 31^ Another study also reported that FAK activation in colorectal cancer cells was attenuated by calcium supplement.^32^ According to these evidences, we supposed that Panx1 reduction in ECFCs may reduce calcium influx, leading to activation of FAK-ERK axis.

To verify the hypothesis, calcium content in non-sense or Panx1 siRNAs transfected ECFCs was evaluated by fluo-4 AM uptake and the results showed that calcium content was reduced by transfection with each of the two Panx1 siRNAs (Figure S11A and S11B). Furthermore, FAK activation was detected in ECFCs treated with BAPTA (calcium intracellular chelator) and showed that phosphorylation of FAK was enhanced with increased concentration of BAPTA (Figure S11C and S11D). In addition, activation of FAK-ERK axis was also examined in Panx1 down-regulated senescent ECFCs supplemented with lactate calcium salt (CaLa). The results showed that activation of FAK-ERK axis induced by Panx1 reduction could be abolished by increased concentration of Cala (Figure 7A to 7C). Furthermore, IGF-1 production improved by Panx1 reduction was also attenuated by Cala (Figure 7D). These findings were summarized in Figure 7E, which showed that calcium influx reduced by Panx1 knockdown activated FAK-ERK axis and then lead to enhanced IGF-1 production to improve angiogenic potential of senescent ECFCs.

**Figure 7.**
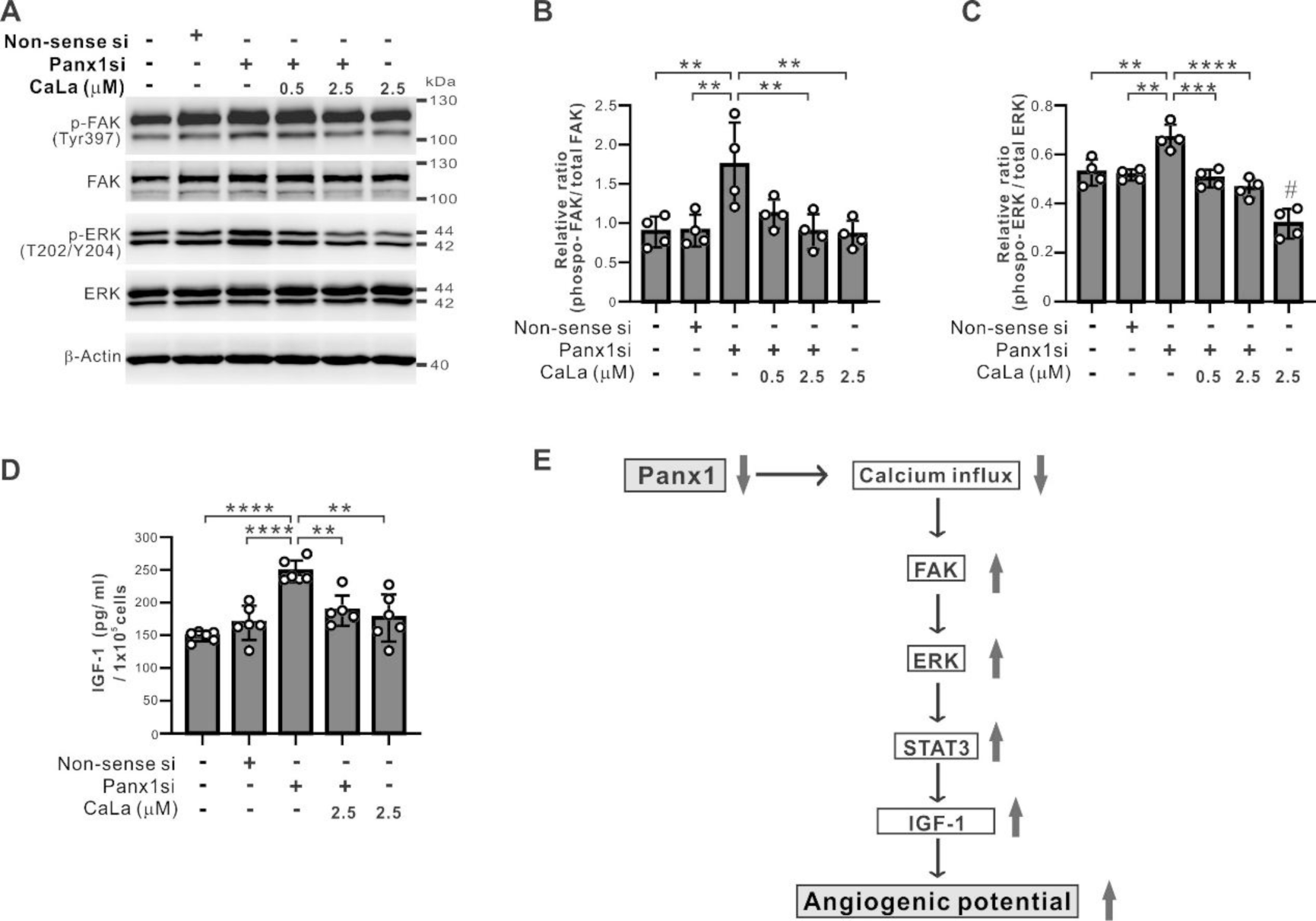
Effects of calcium supplement on FAK-ERK axis activation and IGF-1 production in Panx1 knockdown ECFCs. **A-C**, Western blotting (**A**) and density statistics (**B** and C) showed that enhanced phosphorylation of FAK (**A** and **B**) and ERK (**A** and **C**) in ECFCs at 30 minutes post Panx1 siRNA transfection was abolished by addition of lactate calcium salt (CaLa, 0.5 and 2.5 μM). N=4 ECFC strains from individual donors. **D**, ELISA showed enhanced IGF-1 production in ECFCs after Panx1 siRNA transfection was attenuated by addition of CaLa (2.5 μM). N=6 ECFC strains from individual donors. Data are mean ± SD. Statistical analysis was performed by one-way ANOVA followed by the Tukey’s test. **, P<0.01. ***, P<0.001. ****, P<0.0001. #, P<0.05, compared to each of the other groups. **E,** The working model shows down-regulation of Panx1 reduces calcium influx and then activates FAK, ERK, and STAT3, followed by increased IGF-1 production, leading to enhanced angiogenic potential of senescent ECFCs.

Finally, we also examined eNOS expression in Panx1 overexpressed young ECFCs treated with BATPA due to that calcium influx was related to eNOS production.^33^ The results showed that, in the control young ECFCs, total eNOS expression, but not phospho eNOS ratio, could be increased by calcium influx reduction after BATPA treatment. However, both total eNOS expression and phospho eNOS ratio was not restored by calcium influx reduction in Panx1 overexpressed young ECFCs. These findings indicated that eNOS expression impaired by Panx1 overexpression in ECFCs is through calcium-independent pathway (Figure S12).

## Discussion

Literatures regarding the roles of pannexin isoforms on cellular function and senescence process of EPCs were rare. The present study provided new findings to elucidate its roles as and beyond cell membrane channels of EPCs. First, we confirmed that Panx1 was the predominant isoform existing in pannexin channels of ECFCs derived from circulating EPCs and was markedly increased during senescence *in vitro* and *in vivo*. Second, angiogenic potential of young ECFCs could be impaired by Panx1 overexpression, but enhanced by Panx1 knockdown. Third, cellular activities and senescent status of replication-induced senescent ECFCs could be rejuvenated by Panx1 knockdown through enhancement of IGF-1 production. Fourth, increased production of IGF-1 in the Panx1 knockdown ECFCs resulted from calcium influx reduction with subsequent activation of the FAK-ERK axis. Fifth, decreased blood perfusion in mouse ischemic hindlimbs treated with senescent ECFCs, compared to those treated with young ECFCs, could be improved by Panx1 knockdown in the senescent cells. These findings not only widened the knowledge base of Panx1 but also had clinical implications.

The present study showed that Panx1 expression was up-regulated in replication-induced senescent ECFCs as well as circulating EPCs from aged animals and humans. Previous reports about regulation of Panx1 expression showed that inflammatory cytokine TNF-α could trigger activation of NF-κB pathway to stimulate Panx1 production in endothelial cells.^18^ In parallel, chronic elevation of inflammatory cytokines, including TNF-α, could be found in blood of elderly donors.^22^ These evidences together suggested that up-regulation of Panx1 in circulating EPCs of elderly donors may result from increased inflammatory cytokines in the aged people. Apart from the inflammatory cytokines, ROS, which was induced by replication-induced senescence in endothelial cells may play a role.^34^ ROS was shown to induce opening of pannexin channels in keratinocytes treated with nonmetal haptens.^19^ Panx1 channels and purinergic receptor complex activated by bacteria infection was shown to modulate ATP production to induce ROS generation in gingival epithelial cells.^35^ These evidences indicate that ROS and pannexin channels may interact with each other, leading to up-regulation of pannexin channels in senescent ECFCs. In our study, we provided the evidence that the enhancement of MnSOD in senescent ECFCs was decreased by Panx1 reduction. We also showed that decreased SirT1 of senescent ECFCs was restored by Panx1 reduction. Taken together, these 2 findings suggested that Panx1 channel may mediate ROS production from mitochondria. However, whether ROS could stimulate Panx1 production in replication-induced senescent cells and the underlying mechanism required further experiments.

One important finding in our study is that Panx1 expression level may affect cellular activity of ECFCs. Literatures regarding the effects of Panx1 reduction in vascular disease progression were controversial. In mice with elastase-induced abdominal aortic aneurysms, up-regulation of Panx1 could be found in vascular endothelial cells, and specific knockout of endothelial Panx1 attenuated the aneurysm formation by reducing proinflammatory cytokine expression and immune cell adhesion in aortic tissue.^36^ In contrast, specific knockout of Panx1 in endothelial cells and macrophages in apolipoprotein E-deficient mice may promote atherosclerotic lesions in the aorta.^37^ Our findings in the present study supported that reduced Panx1 expression possessed advantages on ECFC function, based on the results that cellular activities and therapeutic angiogenic potential of senescent ECFCs could be improved by Panx1 knockdown *in vitro* and *in vivo*. Moreover, Panx1 expression was up-regulated in circulating EPCs from elderly donors and overexpression of Panx1 impaired activities of young ECFCs, indicating that the declined function of circulating EPCs in aged subjects resulted from up-regulation of Panx1. These findings implied that Panx1 expression level could be a potential marker for functional evaluation of circulating EPCs and prediction of vascular disease prognosis.

Although protein structure of pannexin and connexin are similar and, in our previous report and the present study, both molecules are up-regulated in replication-induced senescent ECFCs, the two channel proteins expression levels exhibited differential effects on ECFCs. Reduction of Cx43 impaired cellular activities, including proliferation, migration, and tube formation of ECFCs, but reduction of Panx1 improved the activities, as shown in the present study.^20^ These results suggested that enhanced Cx43 expression in senescent ECFCs may exert a compensatory effect to resist senescence, while enhanced Panx1 expression is one of the elements leading ECFCs to senescence. On the other hand, reduction of these two channel proteins in vascular progenitor cells affected different growth factor expression. Reduction of Cx43 attenuated the expression of VEGF in ECFCs and HGF in smooth muscle progenitor cells.^20, 21^ However, reduction of Panx1 increased IGF-1 production in ECFCs. These results indicated that reduction of structurally similar protein channels in vascular progenitor cells or even the same type of cells possess unexpected diverse effects on the cells.

The present study showed that, in mouse hindlimb ischemia model, blood perfusion ratio, limb outcome, and capillary density of mice injected senescent ECFCs with Panx1 knockdown could be improved to the levels close to those of animals injected young ECFCs. Since our previous study of mouse hindlimb ischemia model treated with human ECFCs showed that only a limited number of the ECFCs existed in the ischemic hindlimbs post 14 to 21 days of injection, suggesting that angiogenesis in ischemic tissue enhanced by the injected ECFCs was mainly attributed to the paracrine effects at early stage.^20, 38^ We therefore analyzed supernatants of Panx1 knockdown ECFCs and used antibody neutralization strategy in cellular activity evaluation of senescent ECFCs and HAECs to confirm that IGF-1 masterminded the paracrine effects of Panx1 knockdown ECFCs. Our findings in the present study are consistent with previous reports that IGF-1, by either gene transfer or local administration, could improve neovessel formation in mouse models of cerebral infarction and hindlimb ischemia.^39, 40^ In addition, IGF-1 was suggested to promote ECFCs function through enhancement of VEGF and eNOS production, though the present study showed that Panx1 knockdown-induced IGF-1 production was not associated with eNOS up-regulation.^2^ Furthermore, clinical studies about IGF-1 and ECFC senescence also showed that age-related decline in number, function, and telomerase activities of circulating EPCs can be restored by IGF-1.^41, 42^ These data, together with our findings in the present study, indicated that IGF-1 induced by Panx1 reduction may not only improve angiogenic potential but also rejuvenate EPCs.

The present study also showed that eNOS expression could be impaired by Panx1 overexpression, indicating that ECFC dysfunction caused by Panx1 overexpression involved eNOS down-regulation. Besides, we also found that eNOS production influenced by Panx1 perturbation is not related to calciun influx. The mechanism about how Panx1 up-regulation influences eNOS production needs further clarified.

Regarding FAK-ERK-STAT3 axis mediated IGF-1 production after Panx1 knockdown in senescent ECFCs in the present study, some studies showed that activation of FAK, ERK, or STAT3 (S727) not only promoted cell growth and activities but also resisted senescence. Inhibition of FAK activation could increase SA-β-gal activities through enhancement of senescence pathway related protein expression, such as p53, in cancer cell lines.^43^ ERK activation, reported to be reduced in brain tissue of aged rats, inhibited oxidative stress to improve longevity of mice.^44, 45^ Lacking phosphorylation of STAT3 at serine 727 decreased postnatal survival rate and retarded growth of mice accompanied by low blood IGF-1 concentration.^46^ In addition, other reports about the interaction of these three signal transduction proteins showed that fucoidan may reduce the expression of senescent indicators, such as p21, in ECFCs by improving FAK-ERK axis activation.^3^ STAT3 phosphorylated by ERK at serine 727 could interact with mitochondria complex I to reduce ROS production in cardiomyocytes.^47^ These reports supported our findings in the present study that FAK-ERK-STAT3 axis mediated IGF-1 production was involved in regulation of senescent status in Panx1 knock senescent ECFCs.

Regarding FAK and intracellular calcium, to our knowledge, the present study is the first report that phosphorylation of FAK could be enhanced by decreased intracellular calcium content owing to reduced calcium influx. In endothelial cells, TNF-α treatment was reported to promote calcium influx through increased expression of Panx1 channels to activate NF-κB pathway and then lead to IL-1β production.^18^ The above two findings together implied that calcium influx level could determine the direction of signal transduction pathway to cytokines or growth factors production. Other studies also showed that increased intracellular calcium in airway smooth muscle cells of elderly people and calcium overload in cells may promote ROS production or apoptosis, indicating that reduction of Panx1 may reduce calcium influx in senescent ECFCs to protect the cells from calcium overload-induced injury.^48–50^

One may ask if IGF-1 could be directly applied for therapeutic angiogenesis of ischemic tissues instead of use of ECFCs with Panx1 knockdown, since, in the present study, enhanced angiogenic potential of senescent ECFCs by Panx1 knockdown in hindlimb ischemia mice was through IGF-1. Although IGF-1 had been reported to ameliorate senescent status and enhance angiogenic potential of EPCs, this molecule was known to bind to different receptors, including IGF-1 receptor, IGF-2 receptor, and insulin receptor, leading to increased risk of cancer. Another method was through low dose growth hormone (GH) administration, which enhanced IGF-1 production and EPC number in healthy adults.^51^ Our current study also provided an alternative strategy to induce IGF-1 production in senescent EPCs by Panx1 knockdown. However, whether cell therapy for tissue ischemia using ECFCs with Panx1 knockdown is superior to systemic or local use of growth factor or IGF-1 and avoid the risk of carcinogenesis required further studies. On the other hand, EPCs have been reported to participate angiogenesis in tumor growth, leading to a concern that improved EPC function by reduction of Panx1 may increase the risk of cancer in the body, which needs further verified.

In conclusion, Panx1 expression is up-regulated in human ECFCs with replication-induced senescence and circulating EPCs from elderly donors. Angiogenic potential of senescent ECFCs is improved by Panx1 reduction through increased IGF-1 production via activation of FAK-Erk axis following decreased calcium influx. Our findings expand the roles of Panx1 in biology and provide new strategies to evaluate EPC activities as well as rejuvenate senescent EPCs for cell therapy in therapeutic angiogenesis.

## Acknowledgments

None.

## Highlights

- Panx1 is up-regulated in human ECFCs subject to replication-induced senescence and in circulating EPCs of aged people.
- Panx1 expression level modulates angiogenic potential and senescent status of human ECFCs in vitro.
- Down-regulation of Panx1 rescues therapeutic angiogenic potential of senescent human ECFCs in vitro through enhanced IGF-1 production via activation of FAK-ERK axis following calcium influx reduction.

